# MaxEnt’s parameter configuration and small samples: Are we paying attention to recommendations?

**DOI:** 10.1101/080457

**Authors:** Narkis S. Morales, Ignacio C. Fernándezb, Victoria Baca-Gonzálezd

## Abstract

Environmental niche modeling (ENM) is commonly used to develop probabilistic maps of species distribution. Among available ENM techniques, MaxEnt has become one of the most popular tools for modeling species distribution, with hundreds of peer-reviewed articles published each year. MaxEnt’s popularity is mainly due to the use of a graphical interface and automatic parameter configuration capabilities. However, recent studies have shown that using the default automatic configuration may not be always appropriate because it can produce non-optimal models; particularly when dealing with a small number of species presence points. Thus, the recommendation is to evaluate the best potential combination of parameters (feature classes and regularization multiplier) to select the most appropriate model. In this work we reviewed 244 articles from 142 journals between 2013 and 2015 to assess whether researchers are following recommendations to avoid using the default parameter configuration when dealing with small sample sizes, or if they are using MaxEnt as a “black box tool”. Our results show that in only 16% of analyzed articles authors evaluated best feature classes, in 6.9% evaluated best regularization multipliers, and in a meager 3.7% evaluated simultaneously both parameters before producing the definitive distribution model. These results are worrying, because publications are potentially reporting over-complex or over-simplistic models that can undermine the applicability of their results. Of particular importance are studies used to inform policy making. Therefore, researchers, practitioners, reviewers and editors need to be very judicious when dealing with MaxEnt, particularly when the modelling process is based on small sample sizes.

## Introduction

Environmental niche modeling (ENM), also referred as to predictive habitat distribution modeling (e.g. Guisan & Zimmermann, 2000), or species distribution modeling (e.g. Elith & Leathwick, 2009; Miller, 2010), is a common technique used in a variety of disciplines that use spatial-explicit ecological data, such as landscape ecology (Amici *et al*., 2015), biogeography (Carvalho & Del Lama, 2015), conservation biology (Bernardes *et al*., 2013, Brambilla *et al*., 2013), marine sciences (Bouchet & Meeuwig, 2015; Crafton, 2015), paleontology (Stigall & Brame, 2014), plant ecology (Gelviz-Gelvez *et al*., 2015), public health (Ceccarelli & Rabinovich, 2015) and restoration ecology (Fernandez & Morales, 2016).

The basic principle behind the ENM is the use of environmental information layers and species presence, pseudo-absence and absence points to develop probabilistic maps of distribution suitability (Elith & Leathwick, 2009). Among the available tools for ENM, the maximum entropy approach is one of the most widely used for predicting species distributions (Fitzpatrick *et al*., 2013; Merow *et al.,* 2013). The maximum entropy approach, part of the family of the machine learning methods, is currently available in the software MaxEnt (Phillips *et al*., 2006; https://www.cs.princeton.edu/~schapire/maxent/). MaxEnt can model potential species distributions by using a list of species presence-only locations and a set of environmental variables (e.g. temperature, precipitation, altitude) (Elith *et al*., 2010). Since 2004 the use of MaxEnt has grown exponentially (Figure 1), and nowadays is one of the preferred methods used for predicting potential species distribution among researchers (Merow *et al*., 2013).

**Figure 1.**
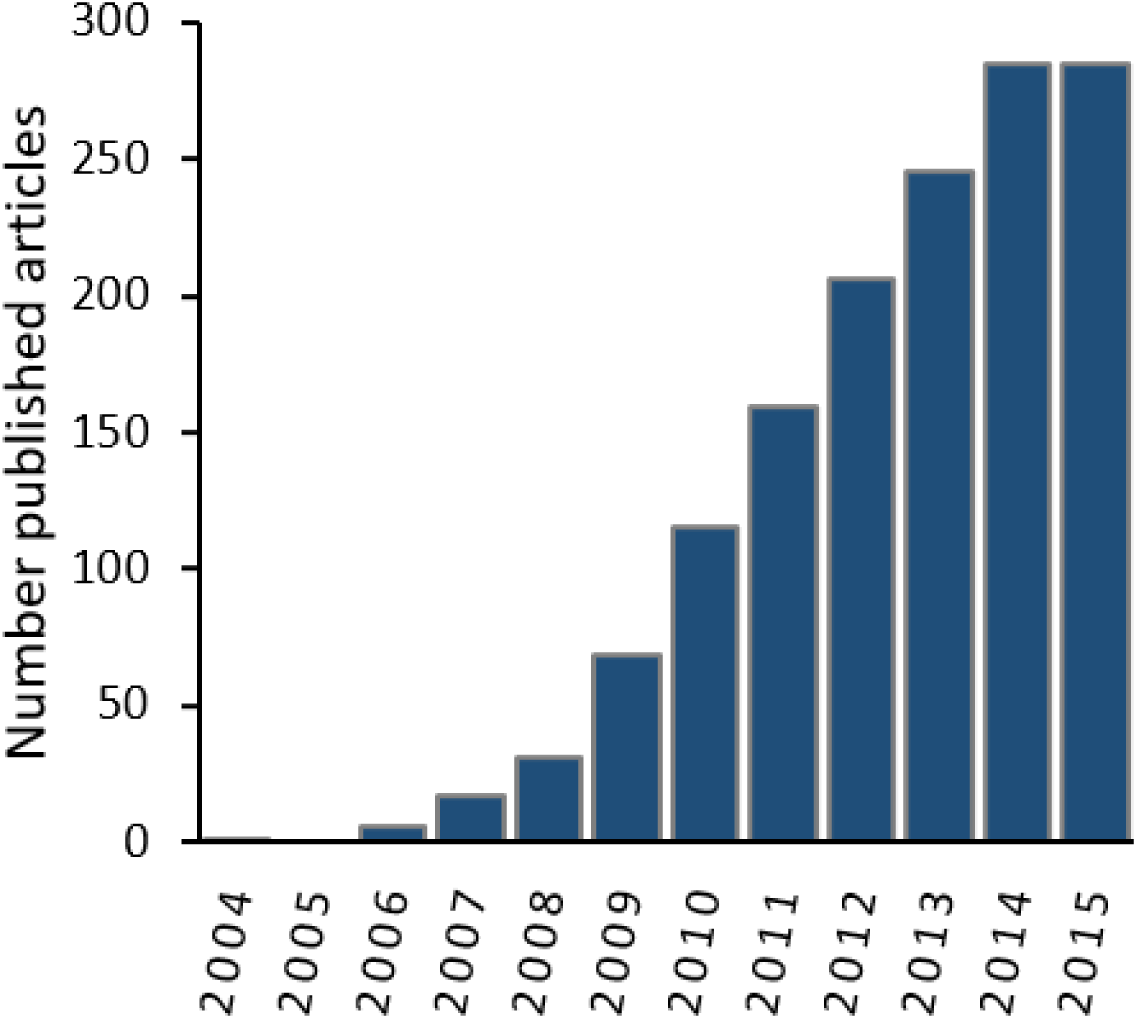
Number of published articles (2004-2015) containing both “MaxEnt” and “species distribution” within the topic in the Web of Knowledge Databases (see methods section for databases details)

The simplicity and straightforward steps required to run MaxEnt seem to have tempted many researchers to use it as a black box despite the increasing evidence that using MaxEnt with default parameter settings (i.e. auto-features) will not necessarily generate the best model (e.g. Shcheglovitova & Anderson, 2013; Syfert *et al*., 2013; Radosavljevic & Anderson, 2014). Some authors have argued that the use of default parameters without providing information on this decision could mean that several of published results could be based in over-complex or over-simplistic models (Warren & Seifert, 2011; Cao *et al*., 2013; Merrow *et al*., 2013). For example, Anderson & Gonzalez (2011) compared different MaxEnt configurations to determine the optimal configuration that minimizes overfitting. Their results showed that in several cases the optimal regularization multiplier was not the default. This is supported by other studies showing that a particular combination of feature classes and regularization multiplier provided better results than the default settings (Syfert *et al*., 2013), and that the default configuration provided by MaxEnt is not necessarily the most appropriate, especially when dealing with small samples size (Warren & Seifert, 2011; Shcheglovitova & Anderson, 2013).

To assess whether researchers are paying attention to recommendations regarding the importance of evaluating the best potential combination of MaxEnt’s parameters for modelling species distribution, in this study we review and analyze the published literature from years 2013 to 2015, focusing our analysis in articles reporting modelling based in small numbers of species presence points (i.e. less than 90 presence points). In addition, we assessed 20 case studies to quantify the potential differences in resulting outputs when using software default parameters instead of analyzing different parameters combinations to identify an alternative best model.

## Literature analysis

We used our own literature search protocol using the databases available through the “web of knowledge” search engine (S1) by using the keywords “MaxEnt” and “species distribution” in the topic (search was done by Morales and Baca-González). Because many of the recommendations were published between 2011 and 2012, we restricted our search to the 2013-2015 period. From the results of this search we only selected studies reporting ≤ 90 presence species points for the modelling process. We chose this threshold value because major changes in MaxEnt auto-features parameters occurs when less than 80 presence records points are used for modelling, implying that a sample of 90 could easily represent less than 80 presence points for modelling due to the required sample points that needs to be set aside for validation purposes. Our preliminary search yielded 816 articles. From these articles, 244 reported a sample size of ≤90 presence points and were therefore used for our analyses (Figure 2, Table 1, see the detailed articles list in S2). We reviewed the methodological information provided in the selected articles to determine the types of feature classes and regularization multiplier used for modelling process. We classified features and regularization multiplier used in each paper in three main categories: (1) user-defined parameters, (2) software default parameters, (3) and no information provided. We also evaluated if the articles provided data on the geographical coordinates of presence points used for the modelling process (i.e. lists of geographical coordinates or species presence maps) necessary for potential replication of the modelling process. We considered only those articles providing information on features, regularization multiplier and geographical coordinates as replicable.

**Figure 2.**
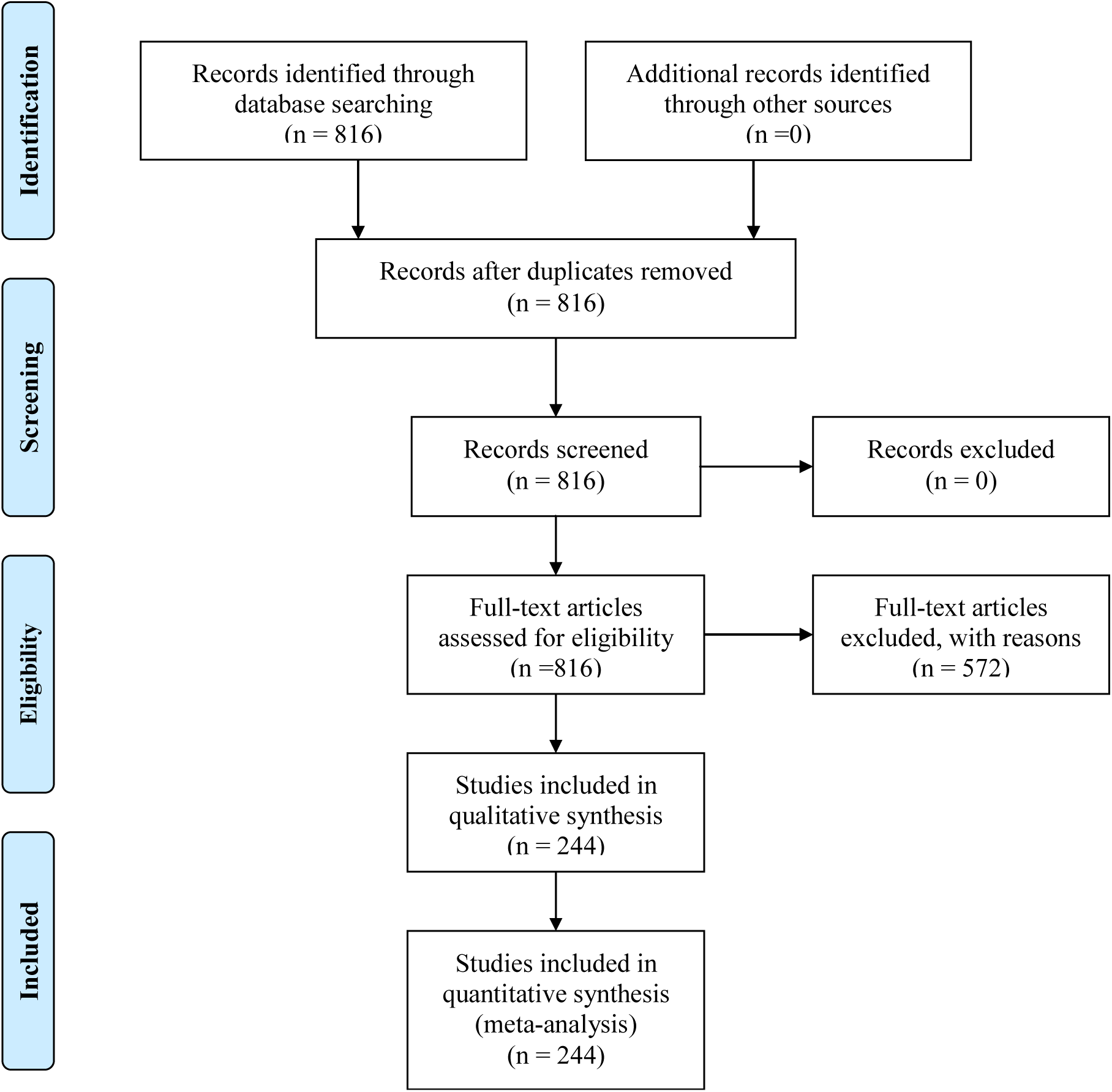
PRISMA flow diagram of the used search protocol following Moher et al. 2009.

**Table 1.**
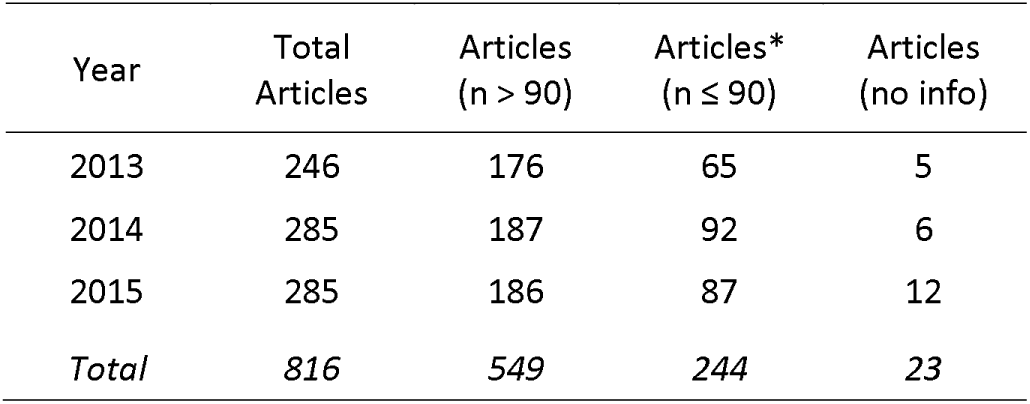
Number of articles published during the years 2013, 2014 and 2015 available through the Web of Knowledge Databases. Articles are presented per year and sample size.

| Year | Total Articles | Articles (n > 90) | Articles* (n ≤ 90) | Articles (no info) |
| --- | --- | --- | --- | --- |
| 2013 | 246 | 176 | 65 | 5 |
| 2014 | 285 | 187 | 92 | 6 |
| 2015 | 285 | 186 | 87 | 12 |
| *Total* | *816* | *549* | *244* | *23* |

## Are we paying attention to recommendations?

Our literature analysis shows that the use of MaxEnt default parameters for modelling species distribution with small recorded presence points seems to be the rule rather than the exception (Figure 3). From the 244 articles that reported a sample size ≤90 for the 2013-2015 period, 44.0% (108 articles) did not provide information about the features used for modelling, 40.0% (97 articles) reported to have used default features, and only 16.0% (39 articles) reported to have used user-defined features (Figure 3; S2). In terms of the regularization multiplier, 48.8% (119 articles) did not provide any information about the regularization multiplier used for modelling, 43.4% (106 articles) used the default regularization multiplier, and only 6.9% (19 articles) reported having used a user-defined regularization multiplier (Figure 3; S2). Considering both default parameters, merely 3.7% (9 articles) of the reviewed articles reported having used user-defined settings for both parameters (S2).

**Figure 3.**
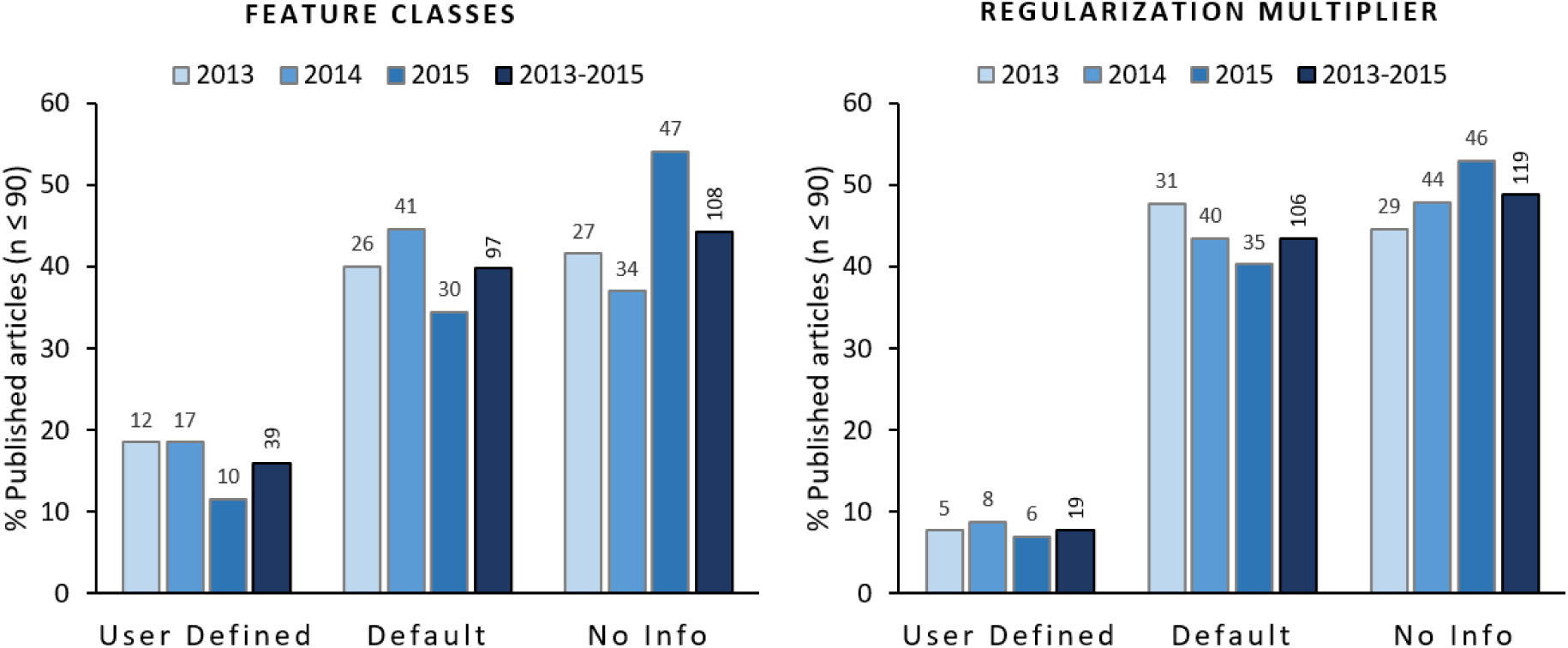
Feature classes and regularization multipliers reported to be used for modelling in the analyzed articles. Columns show the percentage of articles using user-defined, software default, and articles not providing information. Numbers on top of columns represent the number of articles pertaining to each category per year. Columns on the right of each category show the percentage and number of articles for the 2013-2015 period.

## Does ignoring recommendations impact research and practice?

Whereas there is increasing evidence that the use of MaxEnt default parameters do not always generate the best possible model output (e.g. Syfert *et al*., 2013; Radosavljevic & Anderson, 2014), and different authors have highlighted the importance to evaluate the best combination of these parameters before deciding on the best model (see Anderson & Gonzalez, 2011; Warren & Seifert, 2011), results from our analysis indicate that researchers have been rather indifferent to these recommendations. However, the widespread use of default parameters is not the only caveat we found in our literature analysis. We also discovered a general lack of information that would allow the replication or assessment of the results from published studies. In fact, even though 70.5% (172 articles) of publications provide geographical coordinates of presence points, and 47.1% (115 articles) reported both feature classes and regularization multipliers used for modelling; only 34.3% (84 articles) of the analyzed publications provide all three elements together (Figure 4). This information is not only relevant in terms of potential replication of the research, but also necessary for reviewers to evaluate if the outputs from the modelling process are reliable, or are affected among other factors by parameters used, unreliable species presence data sources, or geographically biased presence points records.

**Figure 4.**
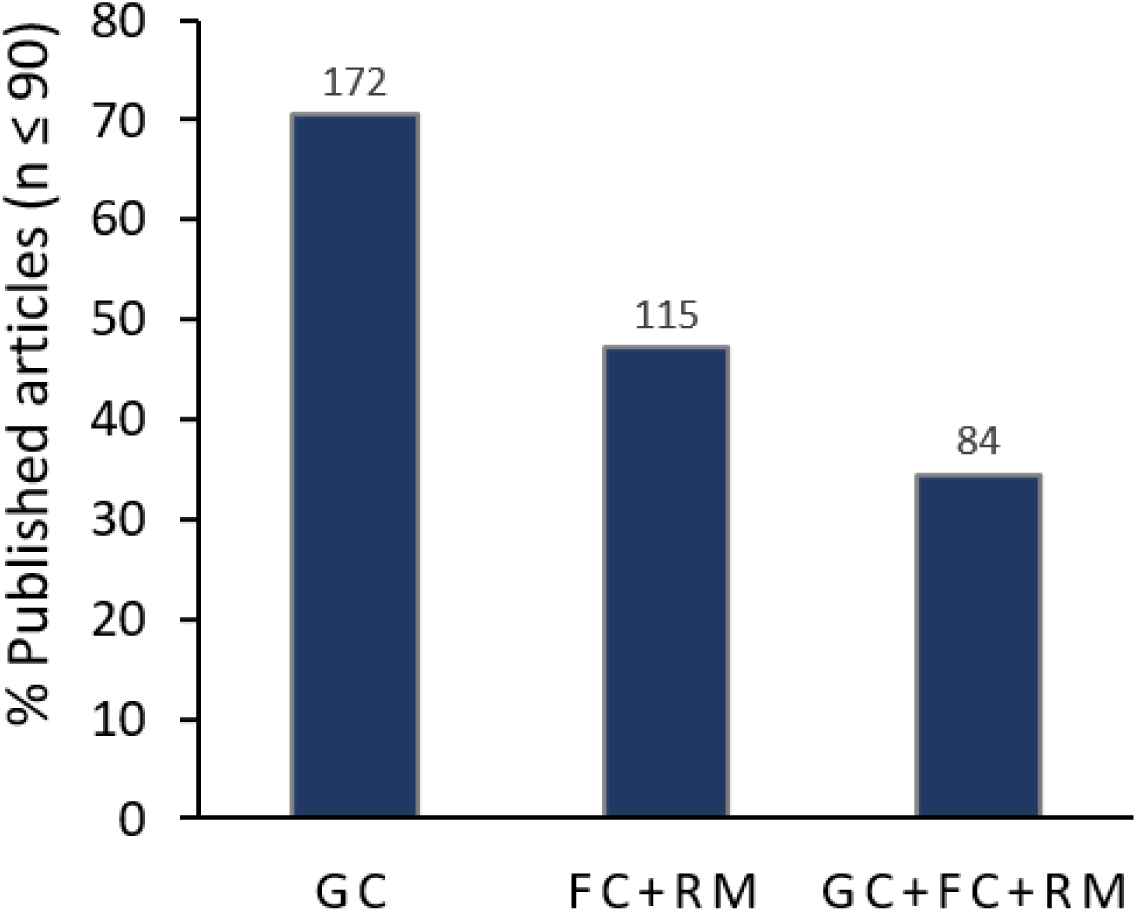
Replicability of the modelling process performed in analyzed articles. Columns show the percentage of articles providing information about GC: geographical coordinates, FC: feature classes, RM: regularization multiplier. Numbers above columns report the number of articles pertaining to each category. Only articles providing information regarding the three inputs (i.e. GC+F+RM column) are considered to provide enough information for replicating the modelling process.

Nevertheless, perhaps the most relevant implications of an inadequate use of MaxEnt for modelling species distribution are on the decision-making arena. When results from the modelling processes are used directly to assess species conservation or to develop conservation strategies, the areas identified as suitable for a given species could differ greatly depending on the parameters using for modelling (Anderson & Gonzalez, 2011). To address this concern, we selected 20 articles from the 84 publications categorized as replicable in our analysis that reported having used default parameters (feature classes and regularization multiplier). We included studies from different regions, with varying geographical extensions, and differing number of species presence points. For each of these articles we collected the geographical coordinates of species presence points and performed the modelling process using default features, and a set of 72 different parameter combinations (See S3), aiming to quantify potential differences on resulting outputs when using default parameters instead of analyzing an alternative best model.

Results from our analysis reveal the huge potential effects of using a default parameter instead of a best model approach for identifying best suitable areas for species distribution (Table 2). Although our results show that the spatial correlation between default and best model outputs is relatively high, and that fuzzy kappa statistics (Visser & Nijs, 2006) show high similarity between generated models for all assessed case studies, the total area identified as suitable for the assessed species tend to greatly differ, particularly for species covering large geographical areas (Table 2). Moreover, it is not only the difference on total area that differs, but also the specific areas that are identified as suitable by both modelling approaches (i.e. shared area). The sample size (i.e. presence points) seems to not affect the differences between the default and the best model outputs, as we did not find a relationship between sample size and models spatial correlation coefficients (R^2^ = 0.026, P = 0.501), fuzzy kappa (R^2^ = 0.005, P = 0.770), or shared/not shared ratio (R^2^ = 0.004, P = 0.786). These results highlight the importance of evaluating what combination of parameters could provide the best modelling results, independently of the sample size used for modelling.

**Table 2.** Estimation of resulting differences when using MaxEnt’s default parameters or a best model approach for modelling species distribution. Spatial correlation values are based in the spatial correlation analysis of MaxEnt’s logistic output. Fuzzy kappa was calculated after applying the 10 percentile training presence logistic threshold to generate the species distribution maps. Area values are based on binary maps generated after applying the 10 percentile training presence logistic threshold.

| Sample Size | Spatial Correlation | Fuzzy Kappa | Area (Km2) |  | Area (Km2) |  | Shared / not Shared ratio | Source |
| --- | --- | --- | --- | --- | --- | --- | --- | --- |
|  |  |  | Default | Best Model | Shared | Not Shared |  |  |
| 7 | 0.856 | 0.864 | 144129 | 447092 | 142612 | 305996 | 0.466 | Carvalho et al. 2015 |
| 8 | 0.957 | 0.799 | 76 | 66 | 66 | 10 | 6.333 | Fois et al. 2015 |
| 9 | 0.905 | 0.797 | 15907 | 9771 | 9212 | 7254 | 1.270 | Chunco et al. 2013 |
| 10 | 0.943 | 0.781 | 861 | 1939 | 843 | 1113 | 0.758 | Alfaro Saiz et al. 2015 |
| 11 | 0.992 | 0.943 | 122415 | 149775 | 121283 | 29624 | 4.094 | Chetan et al. 2014 |
| 12 | 0.983 | 0.841 | 428209 | 551196 | 425674 | 128056 | 3.324 | Palma Perez 2013 |
| 12 | 0.836 | 0.906 | 175166 | 174543 | 156798 | 36113 | 4.342 | Pendersen et al. 2014 |
| 13 | 0.960 | 0.843 | 33421 | 26169 | 24317 | 10957 | 2.219 | Alamgir et al. 2015 |
| 13 | 0.995 | 0.965 | 22013 | 26445 | 21820 | 4818 | 4.528 | Mweya et al. 2013 |
| 14 | 0.948 | 0.916 | 363 | 907 | 353 | 565 | 0.625 | Meyer et al. 2014 |
| 15 | 0.967 | 0.900 | 5004 | 8845 | 4991 | 3867 | 1.291 | Urbani et al. 2015 |
| 16 | 0.769 | 0.652 | 13466 | 28948 | 12848 | 16719 | 0.768 | De Castro et al. 2014 |
| 26 | 0.865 | 0.847 | 5655316 | 7383714 | 5003914 | 3031203 | 1.651 | Chlond et al. 2015 |
| 26 | 0.945 | 0.705 | 32020 | 36420 | 28695 | 11051 | 2.597 | Simo et al. 2014 |
| 31 | 0.937 | 0.879 | 243764 | 248513 | 196113 | 100051 | 1.960 | Orr et al. 2014 |
| 49 | 0.962 | 0.880 | 135239 | 103330 | 100192 | 38186 | 2.624 | Hu et al. 2015 |
| 54 | 0.945 | 0.858 | 2491722 | 1723084 | 1598103 | 1018599 | 1.569 | Confiti et al. 2015 |
| 55 | 0.841 | 0.863 | 1649518 | 1570127 | 1362351 | 494944 | 2.753 | Vergara et al. 2015 |
| 58 | 0.827 | 0.862 | 5822694 | 5370521 | 4439531 | 2314154 | 1.918 | Aguilar et al. 2015 |
| 76 | 0.934 | 0.858 | 3904018 | 3700108 | 3406765 | 790596 | 4.309 | Yu et al. 2014 |

## Implications and future directions

More than 40% of the articles analyzed in our study do not provide information about the parameters configuration used to run the models, which reveals the little attention that researchers and reviewers are paying to this specific issue. Our results also reveal that among the articles that do provide information about the features and regularization multiplier used, a large proportion reported to have used the software default configuration. This preference towards using default setting has remained strong despite the variety of articles describing how MaxEnt works and should be used (Phillips & Dudík, 2008), the proper configuration process (e.g. Merow *et al*., 2013), the potential implications of not selecting the best parameters combination (e.g. Anderson & Gonzalez, 2011; Warren & Seifert, 2011; Syfert *et al*., 2013; Radosavljevic & Anderson, 2014) and the increasing publication of approaches to select the best model by using appropriate parameters combinations (see Anderson & Gonzalez, 2011; Syfert *et al*., 2013; Shcheglovitova & Anderson, 2013).

In addition, we did not observe any trend in the data that would suggest a change from “black box” users towards the use of user-defined parameters. Although our reviewed articles cover a relatively short period of time (2013-2015), if authors were inclined to adopt best practices for modelling we would have expected to see a trend in the data showing an increasing use of user-defined features over time. However, the only clear trend in our results is the increasing number of articles not providing information on the features and regularization multiplier used for modelling. We do not have a clear explanation for this trend, but we believe that it is probably due to new researchers using the modelling software without paying proper attention to current MaxEnt literature, particularly to the publications referring to the importance of analyzing parameters combination for selecting the best model (e.g. Anderson & Gonzalez, 2011; Warren & Seifert, 2011; Syfert *et al*., 2013; Radosavljevic & Anderson, 2014).

## Concluding Remarks

Our results have vast implications, particularly with regard how articles are being reviewed, and the replicability and transferability of the results. We adhere to the calls from other authors to pay better attention to the potential implication of using Maxent’s default parameters when modelling species distribution, but we also suggest reviewers to carefully evaluate if the methodological approach used for modelling is reliable and well supported in recent literature. In addition, researchers need to provide as much information as possible to allow proper evaluation and increase the potential replicability and transferability of their results. These simple recommendations can help to improve the applicability of resulting models, which in turn will help practitioners and decision-makers to use them more effectively as practical tools for the development of management and conservation activities. While the use of MaxEnt’s default parameter can be very useful for having a quick picture of the potential distribution of a given species, taking the necessary time to evaluate what parameters combination results in the best model could largely increase the accuracy and reliability of modelling results.

